# ProteinSketch translates spatial intuition into protein design with bimanual interaction in VR

**DOI:** 10.64898/2026.07.19.739460

**Authors:** Donghyeok Ma, Junyup Lee, Heechan Lee, Sang-Hyun Lee, Yeong Gyeong Oh, Ye-Eun Kim, Taegyu Jin, Siripon Sutthiwanna, Seung-Jun Lee, Woosung Jeon, Dong Sun Lee, Jinsook Ahn, Ryeongeun Cho, Hyemi Park, Seongin Jeong, Hahnbeom Park, Seok-Hyung Bae, Ho Min Kim

**Affiliations:** Department of Industrial Design, Korea Advanced Institute of Science and Technology, Daejeon, 34141, Republic of Korea; Department of Biological Sciences, Korea Advanced Institute of Science and Technology, Daejeon, 34141, Republic of Korea; InnoCORE AI-CRED Institute, Korea Advanced Institute of Science and Technology, Daejeon, 34141, Republic of Korea; Biomedical Research Division, Korea Institute of Science and Technology, Seoul, 02792, Republic of Korea; Department of Biological Sciences, Seoul National University, Seoul, 08826, Republic of Korea; DRB-KAIST SketchTheFuture Research Center, Korea Advanced Institute of Science and Technology, Daejeon, 34141, Republic of Korea

## Abstract

Generative AI–based protein design can create diverse structures for desired functions but offers human designers limited control over three-dimensional architectures. We developed ProteinSketch, a bimanual virtual reality platform in which immersive backbone sketches and volumetric envelopes are directly created and translated into constraints for diffusion-based protein generation. RFdiffusion-based real-time refinement enabled interactive exploration and construction of user-specified protein topologies. Volumetric conditioning of sketched envelopes enabled high-fidelity control of anisometric geometries, confirmed by cryo–electron microscopy. These user-defined volumetric constraints enabled functional-binder design and extension of pre-existing minibinders to surfaces otherwise difficult to access within complex molecular environments using conventional design approaches. This collaborative human–AI framework integrates human spatial reasoning with generative AI for spatially directed, shape-controlled design of protein structure and function.

**One-sentence summary:** Immersive ProteinSketch embeds human spatial reasoning in AI-guided design of protein topology, shape, and function.

## Main Text

Proteins are versatile macromolecular machines whose biological functions are dictated by intricately folded three-dimensional (3D) architectures. Recent breakthroughs in protein AI, encompassing structure prediction (*1, 2*), inverse folding (*3, 4*), and scaffold/structure generation (*5–8*), have revolutionized *de novo* protein design, accelerating applications ranging from precision therapeutics (*9, 10*) to programmable nanomaterials (*11–13*). However, the stochastic nature of computational structure generation and purposeful human design bears a fundamental gap. Although diffusion-based models can generate scaffolds *de novo* or satisfy script-defined constraints such as amino acid length, binding hotspots, and structural symmetry, they do not directly help translate a designer’s spatial intuition into precise structural constraints. Such a limitation is most pronounced when design goals involve complex topologies, target-fitting geometries, or exact spatial envelopes that are intuitive to the human imagination yet difficult to encode through usual text-based or scripting interfaces. Consequently, protein design remains tethered to an expensive “generate-and-hope” cycle of underspecified prompting, massive stochastic sampling, and post-hoc filtering. Although recent approaches have begun to specify protein architectures through top-down structural planning (*14*) or by conditioning generative models on spatial primitives such as curves (*15*) or volumes (*8, 16*), these spatial constraints are often authored indirectly, lack intuitive editing of spatial topologies and geometric constraints, and have limited experimental validation across diverse design objectives.

We reasoned that an immersive interface could bypass the encoding bottlenecks of conventional scripting by enabling designers to translate spatial intuition directly into structural constraints within a native 3D environment. By leveraging virtual reality (VR) and human–computer interaction (HCI) technologies, specifically utilizing intuitive gestures (*17–20*), designers can sketch backbone topologies and define volumetric envelopes that serve as explicit spatial constraints for diffusion-based generative models, such as RFdiffusion, for *de novo* protein design. We, therefore, set out to develop “ProteinSketch,” an interactive VR platform that combines immersive, multiscale sketching with real-time generative AI to integrate hierarchical topology and volumetric-envelope conditioning into a seamless *de novo* protein design workflow.

### ProteinSketch: A bimanual VR platform for translating spatial intent into designable backbones

ProteinSketch is a VR-native, AI-assisted platform that provides a bimanual interactive interface for multiscale intent specification in protein design (**Fig. 1**). Employing bare-hand sketching interactions inspired by natural human coordination (*17–20*), the system allows designers to intuitively specify intricate spatial topologies and geometric constraints in 3D, effectively translating immersive spatial intuition into biophysically viable protein backbones.

**Fig. 1.**
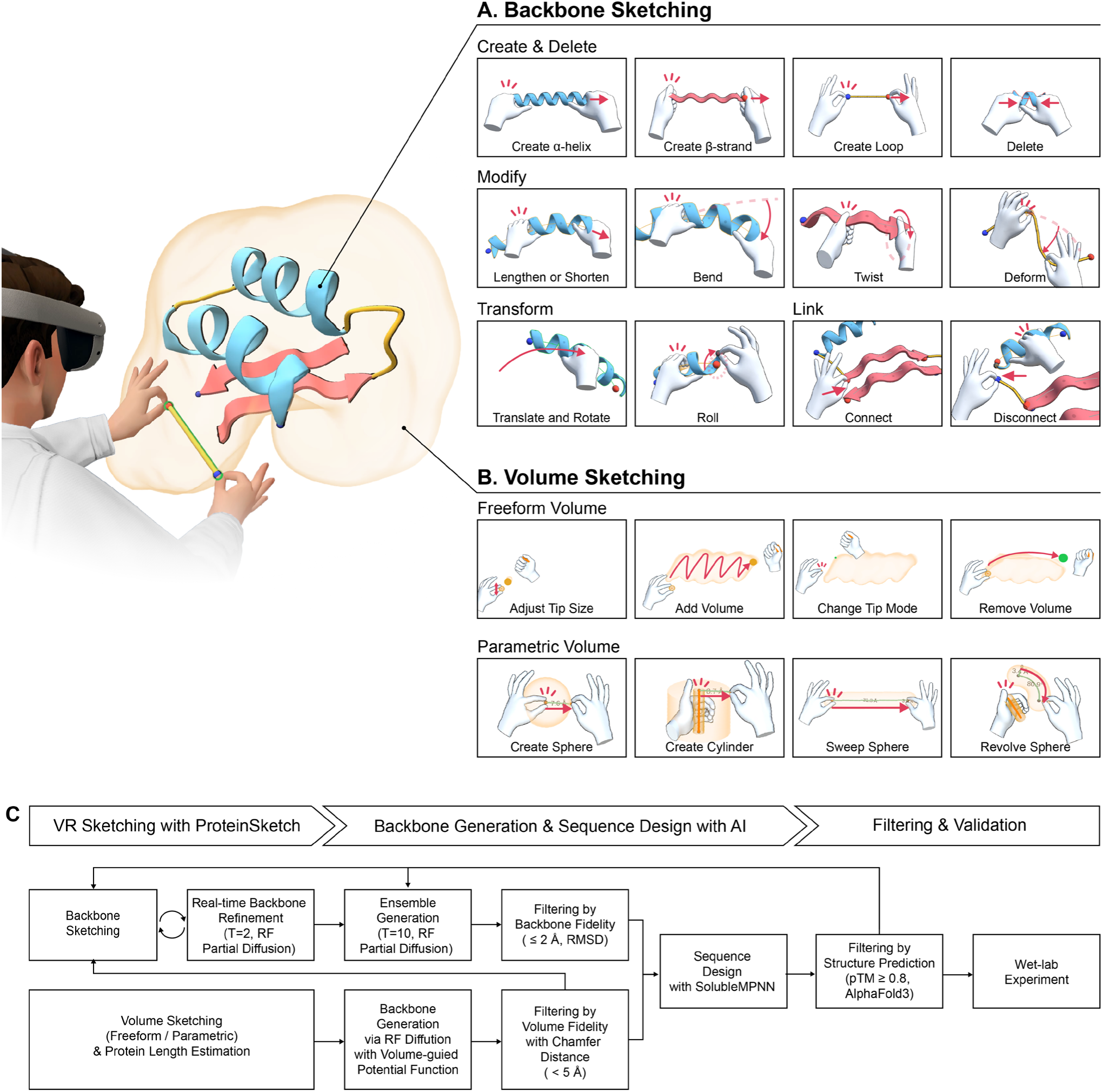
ProteinSketch: immersive VR-native and AI-assisted system for protein design. **(A)** Backbone sketching via bimanual bare-hand gestures to intuitively create and edit secondary-structure elements (α-helices, β-strands, and loops) and assemble them into user-defined topologies. Representative operations are shown: Create & Delete, Modify, Transform, and Link (**see supplementary methods and Fig. S1 for details**). **(B)** Volume sketching for spatial constraint specification. Users define three-dimensional envelopes as either freeform (dynamic addition or removal of density) or parametric (geometry-based profiles) volumes (**see supplementary methods and Fig. S3 for details**). The sketched envelopes served as explicit spatial constraints for conditioned backbone generation. **(C)** End-to-end workflow integrating VR-native “ProteinSketch” with a generative AI–based protein design pipeline. In ProteinSketch, users sketch backbone topology and/or volumetric envelopes. Real-time refinement during iterative backbone sketching and volume-conditioned backbone generation were performed using RFdiffusion (partial diffusion and volume-guided conditional sampling, respectively). Subsequently, sequences were designed with SolubleMPNN, and their structures obtained were filtered with AF2/AF3 applying conventional confidence criteria, followed by wet-lab experimental validation.

First, ProteinSketch enables designers to sketch protein architecture and topology directly within a native 3D environment by placing and editing idealized secondary-structure elements (SSEs) with high spatial precision. It implements a gesture-based control scheme tailored to define explicit SSE parameters, including type, length, position, orientation, curvature, and connectivity (**Figs. 1A and S1**). In particular, α-helices are manipulated via a cylindrical “grab” gesture (analogous to holding a pipe), whereas β-strands

use a planar “thumb-grip” gesture (analogous to holding the edge of a thin plate). During SSE creation, length is specified by anchoring one end with the non-dominant hand while extending the element to the desired terminus with the dominant hand. The resulting SSE can be transformed (translation, rotation, and roll) and structurally modified (lengthening/shortening, bending, and twisting) through intuitive editing gestures and connected via flexible loop segments using a “pinch” gesture (analogous to holding a thin cord) (**Fig. S1**). These interactions follow the principle of asymmetric bimanual coordination (*21*), where the non-dominant hand maintains the spatial frame of reference while the dominant hand performs structural manipulations.

To ensure structural viability during backbone sketching, ProteinSketch couples this SSE-level authoring with real-time structural refinement that corrects local geometric violations within the manual sketch while preserving the intended topological blueprint. This refinement is achieved through the RFdiffusion partial diffusion protocol (**Fig. 1C and Supplementary methods 3.2.1 Real-time refinement of immersive backbone sketches in ProteinSketch**) (*5*). As conventional Rosetta relaxation (e.g., FastRelax) typically requires an explicit all-atom model with side-chain coordinates for physically meaningful refinement, we chose RFdiffusion. In particular, RFdiffusion partial diffusion can operate directly on backbone-only inputs and efficiently regularize local geometry during denoising. This partial diffusion step corrects physiochemical violations introduced by freehand sketching (e.g., chain breaks and severe geometric artifacts) while minimally perturbing the user-specified topology. Consistent with our quantitative benchmarking, major defects are resolved starting at T = 2, which provides the best trade-off between interactive latency and local geometric validity (**Fig. S2 and Supplementary methods 3.2.1 Real-time refinement of immersive backbone sketches in ProteinSketch**). Such an immediate feedback loop enables continuous, iterative “sketch-and-refine” cycles in VR, empowering designers to explore the protein fold space while remaining within the boundaries of designable protein geometries.

Second, ProteinSketch enables designers to create volumetric envelopes that dictate overall protein shape and serve as explicit geometric constraints for diffusion-based backbone generation. While recent advances have introduced structural layout control via 3D ellipsoids (*16*) or 3D curves (*15*), these approaches often require nontrivial parameterization, auxiliary editing, or two-dimensional (2D)-to-3D “lifting” of inputs. ProteinSketch leverages immersive bimanual gestures to directly sculpt volumetric envelopes with native 3D intuition, either as freeform or parametric envelope (**Figs. 1B and S3**). Freeform volumes are well suited for irregular geometries (e.g., fitting into concave binding pockets), whereas parametric ones enable rapid creation and continuous adjustment of analytic 3D shapes.

To translate these sketched envelopes into biophysically viable protein backbones, we developed a volume-guided potential function and integrated it into the RFdiffusion denoising/sampling process to bias backbone trajectories toward occupying the designer-defined envelope (**Fig. 1C and Supplementary methods 3.2.2** (**3**) **Potential functions for volumetric constraints in RFdiffusion**). As volumetric satisfaction is intrinsically coupled to sequence length, we also developed and implemented an envelope-to-length estimation module that converts a sketched volume into an RFdiffusion-ready contig specification, eliminating trial-and-error tuning and enabling rapid end-to-end iteration from envelope sketching to conditioned backbone generation under a consistent geometric constraint (**Supplementary methods 3.2.2** (**2**) **Envelope-to-residue-length estimation**). Computational benchmarking indicates that such volume-guided conditioning outperforms existing voxel-based or ellipsoid-constrained approaches in both designability and structural fidelity to the intended envelope (**Fig. S4**).

Such spatial precision is particularly advantageous for complex binder design, which must satisfy stringent shape complementarity within a prescribed pose while avoiding steric conflicts with adjacent target domains or bound ligands/proteins. By enabling the direct specification of “forbidden” zones and intended “occupancy” regions, ProteinSketch renders backbone generation constraint-directed and designer-steered, substantially reducing trial-and-error and enhancing scaffold placement reproducibility within complex molecular environments. This guided exploration accelerates design iterations for functional binders while maintaining full compatibility with diffusion-based sampling. In both backbone sketching and volumetric conditioning modes, designed backbones proceed through a standard downstream pipeline – sequence design with ProteinMPNN, structural filtering with AlphaFold2 (AF2) and/or AlphaFold3 (AF3), and wet-lab expression and biophysical validation, consistent with established *de novo* protein design workflows (**Fig. 1C**).

### *De novo* protein design through immersive backbone sketching

To evaluate the robustness and generalizability of backbone sketching, we targeted five architecturally distinct *de novo* protein topologies spanning diverse secondary-structure compositions: 2-helix coiled-coils, 3-helix parallel bundles, 3-helix coiled-coils, β-barrels, and β-sandwiches (**Figs. 2 and S5**). For each target, we iteratively sketched SSEs (α-helices and β-strands) and refined their geometry within the ProteinSketch VR environment using RFdiffusion (partial diffusion, T = 2) to specify their spatial orientations, alignments, and loop connectivity (**Figs. 2A and S2**).

**Fig. 2.**
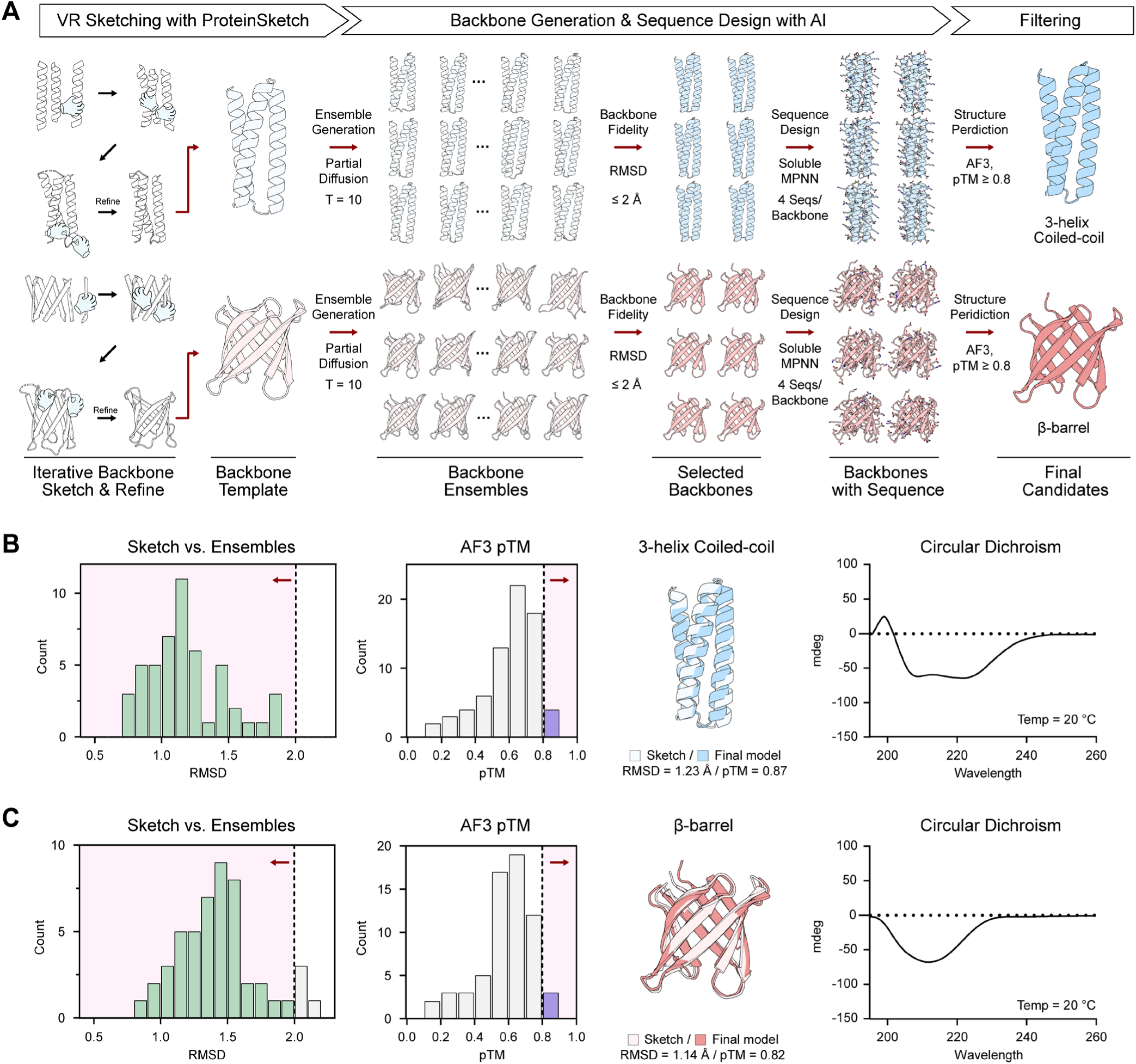
*De novo* protein design through immersive backbone sketching with ProteinSketch. **(A)** Overview of the “backbone-sketch-to-design” workflow for representative targets: 3-helix coiled-coils and β-barrels. Iterative VR sketching and refinement with ProteinSketch produced a user-defined backbone template, which was expanded into backbone ensembles by RFdiffusion partial diffusion (T = 10). Ensemble members were filtered by backbone fidelity to the sketch (RMSD ≤ 2 Å), followed by sequence design with SolubleMPNN. Designs were sequentially filtered with AF2 in initial-guess mode (backbone RMSD ≤ 1.0 Å, pLDDT ≥ 85, and pAE ≤ 10), followed by AF3 prediction (pTM ≥ 0.8) as the final selection criterion. **(B and C)** Design and experimental characterization of the 3-helix coiled-coil **(B)** and β-barrel **(C)**. Left to right: Distributions of sketch-to-ensemble RMSD and AF3 pTM scores with filtering thresholds indicated (dashed lines). A representative final design and structural comparison (Gray, sketched model; Colored, AF3 prediction). Circular dichroism spectra acquired at 20°C confirm the intended secondary-structure composition. Additional designs (2-helix coiled-coil, 3-helix parallel bundle, and β-sandwich) are provided in **Fig. S5**.

Although interactive sketch–time refinement uses short denoising (T = 2) to provide real-time feedback while preserving the sketched topology, we subsequently applied RFdiffusion to the sketched backbone template with longer denoising (partial diffusion, T = 10) to convert the coarse manual sketch into a diverse ensemble of designable backbones for downstream sequence exploration. This timestep can be adjusted per the design objectives: lower values preserve sketch fidelity for higher-confidence designs, whereas greater values facilitate broader conformational exploration for increased backbone diversity. This stage serves as a critical structural refiner, relaxing gestural inputs into biophysically favorable ensembles while preserving the intended global topology. For the 3-helix coiled-coil, the resulting ensemble exhibited high architectural fidelity to the initial VR sketch (mean RMSD = 1.2 Å) (**Fig. 2B**). In contrast, the β-barrel design required more substantial geometric regularization (mean RMSD = 1.6 Å) to rectify intricate inter-strand hydrogen-bonded networks that are difficult to define manually with sub-angstrom precision (**Fig. 2C**).

We selected ensemble members with RMSD ≤ 2.0 Å to preserve global fold fidelity while maximizing backbone diversity for sequence exploration (**Figs. 2B and 2C**). Following sequence design with SolubleMPNN (*22*), candidates were evaluated via AF2 in the “initial-guess” mode (*1, 23*). Designs were filtered based on the following criteria: backbone RMSD ≤ 1 Å, pLDDT ≥ 85, and pAE ≤ 10. Candidates satisfying these criteria were subsequently validated using AF3 (*2*), where they were retained only if they achieved pTM ≥ 0.8 and pLDDT ≥ 85. Only designs that passed both filtering were prioritized for experimental validation. Representative designs satisfied geometric fidelity and confidence criteria: for example, sketch-to-model RMSD = 1.23 Å with pTM = 0.87 for the 3-helix coiled-coil; and RMSD = 1.14 Å with pTM = 0.82 for the β-barrel (**Figs. 2B and 2C**).

Experimental characterization corroborated the high success rate of this integrated pipeline. Across topology classes, 75%–90% of synthesized variants were expressed as soluble, monodispersed proteins in *Escherichia coli*, as verified by size-exclusion chromatography (**Fig. S5**). Circular dichroism spectroscopy further supported the intended secondary-structure compositions (**Figs. 2B, 2C, and S5A–S5C**). We benchmarked using TopoBuilder, which use string-based descriptors defining SSEs (*24, 25*). It requires iterative manual tuning to approximate the hand-sketched 3D geometry, yet it produces consistently low TM-scores, indicating a limited ability to capture user intent and a time-consuming workflow (**Fig. S6**). Taken together, immersive backbone sketching with ProteinSketch provides an efficient, generalizable framework for rapid exploration of *de novo* protein fold space guided by human spatial intuition.

### Volumetric conditioning for high-fidelity shape control in protein design with sketched envelopes

Next, to assess the robustness and generalizability of volumetric conditioning, we tested shape-controlled generation across four geometric classes (spheres, rods, curved arcss, and cones), spanning 100–500 residues (**Fig. 3**). Utilizing the parametric envelope tools in ProteinSketch, we generated the corresponding volumetric envelopes, with target protein lengths determined by employing the envelope-to-length estimation module described above. These sketched envelopes were subsequently encoded as signed distance function (SDF)-based voxel representations (*26*) that provide the spatial field used to compute the conditioning potential during RFdiffusion sampling (**Fig. S7A and Supplementary methods 3.2.2** (**1**) **Converting user-sketched envelope to SDF voxel representation**).

**Fig. 3.**
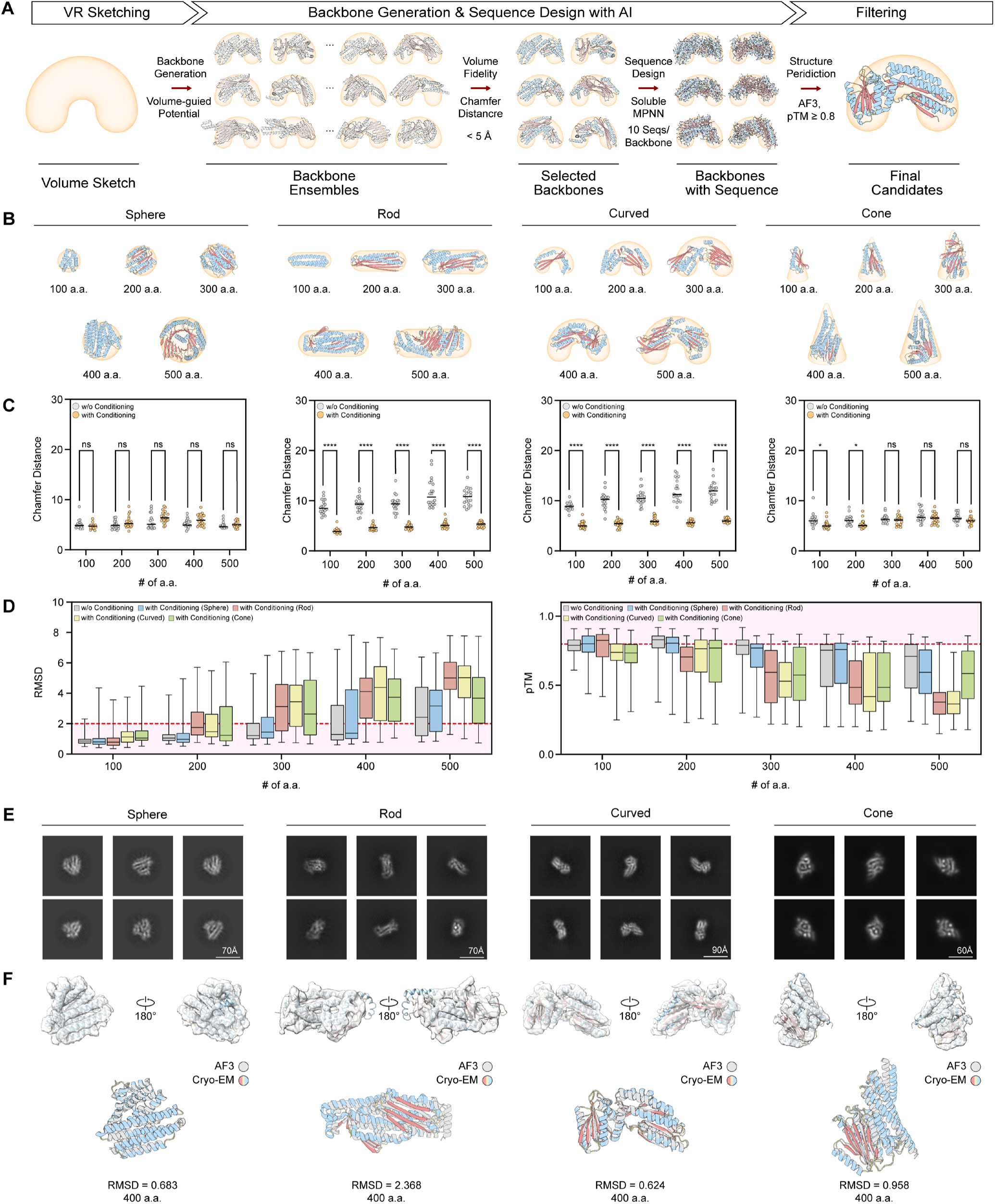
*De novo* protein design through volumetric conditioning with sketched envelopes. **(A)** Overview of the “volume-sketch-to-design” workflow. A volumetric envelope drawn in ProteinSketch was converted into an SDF-based voxel representation, from which a volume-guided conditioning potential was computed during RFdiffusion denoising (**see Supplementary methods 3.2.2** (**3**) **Potential functions for volumetric constraints in RFdiffusion for details**). The backbone ensembles generated were filtered by shape fidelity using Chamfer distance (<5 Å), followed by sequence design with SolubleMPNN (10 sequences per backbone) and structure-based filtering/prioritization using AF2 initial-guess mode: backbone RMSD ≤ 2.0 Å, pLDDT ≥ 85, pAE ≤ 10; and AF3, pTM ≥ 0.8. **(B)** Representative volume-conditioned backbone designs for four target geometries (sphere, rod, curved arc, and cone) across increasing target lengths of 100–500 amino acids, shown within the corresponding volumetric envelopes. **(C)** Shape fidelity quantified based on the Chamfer distance between voxelized designed backbones and the corresponding target envelopes for designs generated with (gold) and without (gray) volumetric conditioning: n = 20 backbones per geometry and target length. Statistical significance was assessed using multiple t-tests. ns, not significant; *P < 0.05; ****P < 0.0001. **(D)** Designability assessed via structural agreement and confidence, shown as backbone RMSD between RFdiffusion backbones and their corresponding AF2-predicted models (left), together with AF3 pTM distributions (right). n = 100 designs (20 backbones × 5 sequences per backbone) per geometry and residue lengths of 100, 200, 300, 400, and 500 amino acids. Red dashed lines indicate the designability thresholds: backbone RMSD ≤ 2 Å and AF3 pTM ≥ 0.8. **(E)** Representative cryo-EM 2D class averages of selected designs for each geometry, ∼400 residues long, exhibiting particle morphologies highly consistent with the intended global shapes. Scale bars are indicated. **(F)** Cryo-EM density maps and refined atomic models (top). AF3-predicted models (gray) overlaid with the corresponding cryo-EM structures (colored), showing close structural agreement (bottom). Backbone RMSD values are indicated.

After generating backbone ensembles with RFdiffusion under volumetric conditioning using the corresponding SDF voxel representation and guiding potential function (**Figs. S7B and S8**), we assessed geometric fidelity by voxelizing the backbones obtained and calculating Chamfer distances to the original sketched envelopes (**Fig. S9 and Supplementary methods 3.2.2** (**4**) **Shape fidelity metric—Chamfer distance and 3.3.2. Assessment of shape fidelity and designability for volume-conditioned monomer design**). Notably, these analyses revealed a clear trade-off between shape fidelity and designability. In particular, a substantially larger fraction of designs achieved low Chamfer distances to the target envelope, indicating that explicit conditioning enforced strict adherence to the sketched volumes (**Fig. 3C**). However, this stronger geometric enforcement was accompanied by increased backbone RMSD and caused a length-dependent decline in AF3 pTM (**Fig. 3D**). Consequently, as the amino acid length increased, only a subset of candidates satisfied the designability criteria: RMSD ≤ 2Å and pTM ≥ 0.8, suggesting that improved conformity to the predefined envelope can compromise foldability and designability.

Interestingly, volumetric conditioning proved to be more beneficial in anisometric geometries (**Fig. 3C**). Spheres exhibited low Chamfer distances even without volumetric conditioning, reflecting an inherent bias of diffusion-based models toward compact, globular architectures. In contrast, for rods and curved arcs, volumetric conditioning markedly reduced Chamfer distances relative to unconditioned sampling across all lengths, indicating that explicit volume-guided potentials are indispensable for translating these anisometric envelopes into viable protein geometries (**Fig. 3C**). For experimental validation, we selected representative designs of ∼400 residues from each geometry class (**Fig. S10**).

To overcome the reduced designability observed in anisometric geometries (rods, curved arcs, and cones), we performed an additional RFdiffusion partial diffusion and refinement (*27*) with selected first-round conditioned backbones as templates. Such an approach expanded the pool of candidates satisfying the structure-prediction confidence criteria (**Figs. S10B–S10D**). Backbones with high geometric fidelity (Chamfer distance < 5 Å) were subsequently processed for sequence design using SolubleMPNN (10 sequences per backbone). The resulting designs were filtered via AF2 (initial-guess mode, backbone RMSD ≤ 2 Å, pLDDT ≥ 85, and pAE ≤ 10) and further prioritized with AF3 (pTM ≥ 0.8 and pLDDT ≥ 85) for final experimental characterization.

The candidate designs were expressed in *E. coli*, purified as monodisperse proteins, and their structures were determined by cryo-electron microscopy (cryo-EM) (**Fig. S11**). The resulting 2D class averages revealed particle morphologies highly consistent with designer-specified envelopes across all geometries (**Fig. 3E**). Subsequent 3D reconstructions and structural refinements demonstrated atomic-level agreement with the design models: RMSD = 0.7–2.4 Å, cryo-EM structure relative to AF3-based predictions (**Fig. 3F**). These validated that ProteinSketch can translate immersive volumetric intuition into high-fidelity protein architectures with near-atomic precision. Collectively, these results demonstrated that the ProteinSketch workflow overcame the stochastic sampling biases plaguing generative models to achieve reliable spatial control over protein morphology.

### Volumetric conditioning for functional-binder design within complex molecular environments

We next asked whether volumetric conditioning could improve binder design efficiency within complex molecular environments. In contrast to the volume-conditioned monomer design described above, high-affinity binder design is fundamentally constrained by the requirement for interface complementarity, including both the shape and chemical properties of the target surface (*28*). Furthermore, to develop functional modulators, such as antagonists or agonists, designers must intentionally impose occupancy regions to engage functionally important sites and forbidden zones to avoid steric conflicts with the target, ensuring that productive sampling satisfies not only interface complementarity but also precise spatial placement within a narrow predefined region. Notably, conventional hotspot-guided RFdiffusion tends to generate elongated helical bundles rather than structurally compact, globular binders. By allowing designers to directly define exclusion and occupancy constraints while enforcing structural compactness, ProteinSketch transforms backbone generation into a constraint-directed, designer-steered process. We reasoned that this framework would reduce the trial-and-error drawback inherent to stochastic sampling and facilitate the generation of compact, site-specific binders tailored to complex target interfaces.

We first applied volumetric conditioning to *de novo* binder design against MD2, a TLR4 co-receptor that mediates lipopolysaccharide (LPS)-induced innate immune signaling and represents a therapeutic target for sepsis intervention (*29–31*). Structurally, MD2 adopts a β-cup fold containing a deep hydrophobic pocket, whereas its pocket entrance rim is enriched in positively charged residues that engage the negatively charged phosphate groups of LPS (**Fig. 4A**). Rather than designing a globally hydrophobic scaffold to occupy this pocket in a manner analogous to the acyl chains of LPS, which could compromise solubility and promote nonspecific aggregation, we sought to create a compact binder in which a hydrophobic segment limitedly inserts into the pocket while the remainder of the scaffold engages the charged pocket rim of MD2 (**Fig. 4B**). However, hotspot-guided RFdiffusion using Leu87, Pro88, and Val93 as constraints, without volumetric conditioning, did not reliably enrich compact backbones at the intended pocket-proximal site and frequently produced poorly packed or mispositioned interfaces (**Figs. 4C and 4D, left**).

**Fig. 4.**
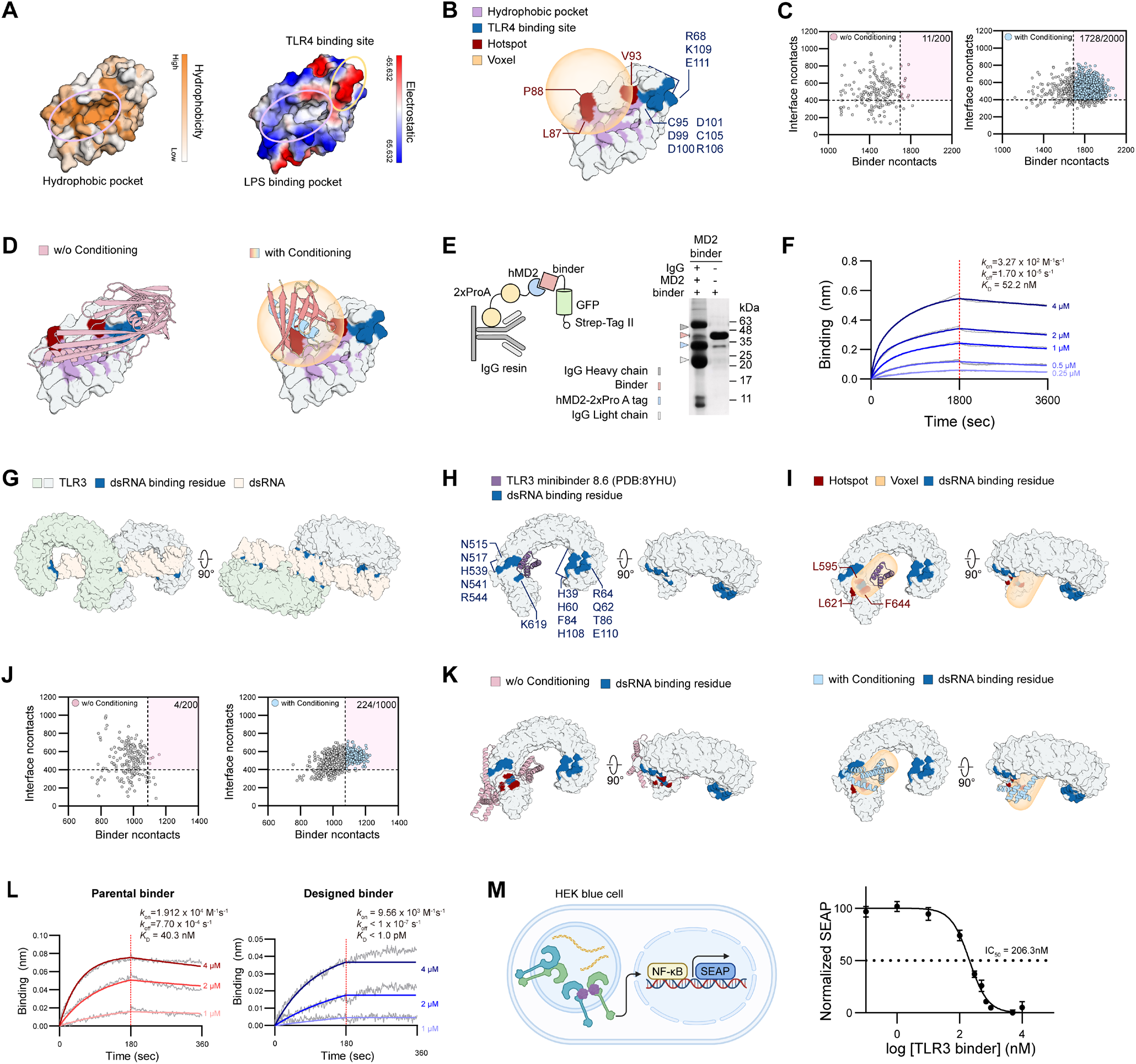
Functional-binder design within complex molecular environments through volumetric conditioning. **(A)** Surface representations of MD2 colored by hydrophobicity (left) or electrostatic potential (right), highlighting the hydrophobic pocket and LPS-binding site (PDB: 3FXI). **(B)** Rationale for volume-conditioned MD2 binder design. The hydrophobic pocket, TLR4-binding site, designated hotspot residues (Leu87, Pro88, and Val93), and the sketched volumetric envelope are shown. **(C)** Contact-based compactness analysis of MD2 binder backbones generated without (left) or with (right) volumetric conditioning. Ncontacts scores of MD2–binder interface contacts (Interface ncontacts) and ncontacts scores of intra-binder contacts (Binder ncontacts) for individual backbones were plotted. Dashed lines indicate applied thresholds (interface ncontact ≥ 400; binder ncontact ≥ 1,700), and values in the upper-right corners indicate the number of backbones satisfying both criteria. **(D)** Representative binder backbones generated employing hotspot-guided RFdiffusion without (left) or with (right) volumetric conditioning. The volume-conditioned backbone occupied the prescribed pocket-proximal region with a compact conformation. **(E)** Pull-down assay for selected MD2 binder candidates. Left: schematic of the pull-down strategy using IgG resin and 2×ProA-tagged MD2 to capture Strep-tag II–GFP-fused binders. Right, SDS-PAGE analysis of the indicated pull-down samples. **(F)** Bio-layer interferometry (BLI) sensorgrams for the selected MD2 binder at the indicated concentrations, with kinetic parameters obtained by global fitting indicated. Red dashed lines represent the transition from association to dissociation. **(G)** Structure of the human TLR3 ectodomain bound to dsRNA, with dsRNA-binding residues highlighted in blue (PDB: 7WV5). **(H)** Binding mode of the parental TLR3 minibinder 8.6 (purple; PDB: 8YHU) onto the concave surface of TLR3. The adjacent dsRNA-binding residues are presented in blue. **(I)** Rationale for volume-guided extension of the parental TLR3 minibinder. The composite volumetric envelope defining the intended extension region is shown in orange, with designated hotspot residues (Leu595, Leu621, and Phe644) highlighted in red. **(J)** Contact-based compactness analysis of TLR3 binder-extension backbones generated without (left) or with (right) volumetric conditioning. Ncontacts scores of TLR3–binder interface contacts (Interface ncontacts) and ncontacts scores of intra-binder contacts (Binder ncontacts) for individual backbones were plotted. Dashed lines indicate applied thresholds (Interface ncontact ≥ 500; Binder ncontact ≥ 1,100), and values in the upper-right corners represent the number of backbones satisfying both criteria. **(K)** Representative extended binder backbones generated utilizing RFdiffusion-based scaffold extension without (left) or with (right) volumetric conditioning. The volume-conditioned design formed a compact extension that occupied the prescribed ligand-binding region. **(L)** BLI sensorgrams comparing the parental TLR3 minibinder (left) and the selected extended binder (right) at the concentrations indicated. Red dashed lines represent the transition from association to dissociation. Kinetic parameters obtained by global fitting are shown. **(M)** Functional characterization of the selected TLR3 binder extension in TLR3-overexpressing HEK-Blue reporter cells. Left: schematic of the NF-κB-dependent secreted embryonic alkaline phosphatase (SEAP) reporter assay. Right: dose-dependent inhibition of poly(I:C)-induced TLR3 signaling by the extended binder. Data represent mean ± SEM from two independent experiments performed in triplicate. Dose–response curves were fitted with a four-parameter variable slope model in GraphPad Prism, and the fitted IC_50_ is shown.

To directly impose the desired binding geometry, we used ProteinSketch to create a volumetric envelope adjacent to the MD2 pocket entrance and combined this spatial constraint with hotspot residues (**Fig. 4B**). The sketched envelope was converted into an SDF-based voxel representation, from which a volume-conditioning potential was computed during RFdiffusion denoising to direct backbone trajectories toward the prescribed pocket-proximal region. In parallel, hotspot constraints at Leu87, Pro88, and Val93 guided interface formation toward the MD2 pocket rim. Together, compared to hotspot-guided sampling alone, these volume- and hotspot-guided constraints substantially enhanced target-proximal binder contacts and enriched compact-backbone conformations occupying the prescribed region (**Figs. 4C and 4D, right**).

Sequences were designed with SolubleMPNN for the selected volume-conditioned backbones (**highlighted by blue circles in Fig. 4C, right**) and initially screened using MSA-free AF3. Candidates geometrically agreeing with the intended target-proximal placement were then re-evaluated using MSA-enabled AF3 (**Fig. S12A**). A chosen first-round backbone was subsequently refined by RFdiffusion (partial diffusion, T = 20), followed by a second round of SolubleMPNN sequence design and structure-based filtering using AF2 in initial-guess mode and MSA-enabled AF3 (**Fig. S12B**). Five final candidates were expressed, purified, and evaluated for MD2 binding by pull-down assay, of which one emitted a positive binding signal (**Figs. 4E, S12C, and S12D**). Bio-layer interferometry further confirmed the direct binding of this design to MD2 with nanomolar affinity: *k*_on_ = 3.27 × 10^2^ M^−1^ s^−1^, *k*_off_ = 1.7 × 10^−5^ s^−1^, and *K*_D_ = 52.2 nM (**Fig. 4F**). The predicted complex structure recapitulated the intended engagement of the MD2 hydrophobic pocket, with well-packed interactions around the pocket entrance rim (**Fig. S12E**). Together, these findings suggest that volumetric conditioning can redirect diffusion sampling toward spatially restricted target sites that remain inefficiently accessed by conventional hotspot-guided design.

We next addressed a more demanding binder-extension problem. We previously designed a high-affinity *de novo* minibinders targeting the concave surface of the human TLR3 and further engineered multivalent variants that promote TLR3 oligomerization and downstream signal activation (*32*). However, the parental minibinders engaged a site distinct from the ligand-binding surface for dsRNA or poly(I:C), thereby limiting their utility as antagonists (*33–35*) (**Figs. 4G and 4H**). Therefore, we sought to extend the parental scaffold over this adjacent ligand-binding surface of TLR3 while preserving its original interaction mode (**Fig. 4I**). Conventional RFdiffusion-based scaffold extension frequently generated elongated, but structurally poorly defined modules that failed to adequately and precisely cover the intended functional surface precisely, underscoring the difficulty of appending a functionally competent extension to a pre-existing binder scaffold. To address this limitation, we used ProteinSketch to create a composite volumetric envelope spanning both the parental binder footprint and the intended extension region overlying the TLR3 ligand-binding surface (**Fig. 4I**). The sketched composite envelope was converted into an SDF-based voxel representation and used during RFdiffusion denoising to guide the extension of the backbone produced toward the prescribed region. In parallel, the hotspot residues Leu595, Leu621, and Phe644 directed interface formation toward the ligand-binding surface. Compared with conventional RFdiffusion-based scaffold extension, such combined volume- and hotspot-guided conditioning yielded a markedly greater fraction of extended binder backbones that satisfied positional occupancy and structural compactness criteria (**Figs. 4J and 4K**). Sequences were designed with SolubleMPNN for backbones passing these filters (**highlighted by blue circles in Fig. 4J, right**) and evaluated with AF2 in initial-guess mode. Candidates were subsequently evaluated utilizing MSA-enabled AF3 and prioritized by manually inspecting interface packing and spatial coverage (**Figs. S12F–S12H**).

Two selected candidates (**Figs. S12F and S12G)** were advanced for experimental characterization. Both were readily expressed and bound with TLR3, but candidate 1 bound more strongly than the parental minibinder and was therefore functionally evaluated (**Figs. S12I and J**). Compared with the parental minibinder (*k*_on_ = 1.91 × 10^4^ M^−1^ s^−1^, *k*_off_ = 7.70 × 10^−4^ s^−1^, and *K*_D_ = 40.3 nM), candidate 1 had a modestly reduced association rate (*k*_on_ = 9.56 × 10^3^ M^−1^ s^−1^), potentially reflecting the greater entropic cost of encounter complexation across a larger interactive surface (**Fig. 4L, left**). In contrast, its dissociation rate was markedly reduced (*k*_off_ < 1 × 10⁻^7^ s^−1^), substantially prolonging complex lifetime and yielding an apparent affinity of *K*_D_ < 1 pM (**Fig. 4L, right**). Consistent with these kinetic parameters, the predicted complex structure indicated that the designed extension preserved the parental binding footprint while establishing additional contacts across the adjacent ligand-binding surface, thereby expanding the overall binding interface (**Fig. S12K**). We then tested whether the designed extension covering the ligand-binding surface conferred functional antagonism. In TLR3-overexpressing HEK-Blue reporter cells, candidate 1 inhibited poly(I:C)-induced TLR3 signaling in a dose-dependent manner, with an IC_50_ of 206.3 nM (**Fig. 4M**). These results demonstrate that volume-guided scaffold extension can effectively translate biological and structural intuition into designable, high-fidelity functional modulator scaffolds by enabling the intended expansion of interfacial contact area and structural complementarity.

## Discussion

We demonstrate that integrating human spatial intuition with generative AI can transform protein design from a largely stochastic sampling task toward a more constraint-directed engineering process. Although current state-of-the-art methods, such as RFdiffusion, Chroma, and BindCraft (*5, 7, 8*), are remarkably versatile for designing *de novo* proteins and binders, they often rely on post hoc selection from large stochastic design pools and generally lack the explicit spatial guidance required to precisely place structural motifs and enforce intended geometry within complex molecular environments. ProteinSketch addresses these limitations by enabling designers to directly dictate topology and overall shape within a native 3D environment, thereby translating immersive geometric reasoning into explicit constraints for backbone generation. Consequently, our most critical contribution is establishing a collaborative human–AI framework that links intuitive spatial reasoning to designable protein architectures. Moreover, by coupling bimanual sketching with real-time diffusion-based regularization, the system facilitates an interactive exploration of protein fold space while ensuring physically plausible backbone geometry. Such a designer-steered approach further enables the direct specification of occupancy regions and forbidden zones, effectively narrowing the search space toward high-fidelity candidates that satisfy predefined geometric and functional intents.

Beyond these conceptual advances in protein design, our results demonstrate that ProteinSketch is applicable across multiple scales of protein design. At the fold level, immersive backbone sketching coupled with real-time diffusion-based regularization enabled the generation of structurally diverse topologies, including helical bundles, β-barrels, and β-sandwiches, but also facilitates the direct interactive editing of protein architectures within a native 3D environment. At the shape level, volumetric conditioning enabled high-fidelity generation of proteins matching prescribed envelopes, particularly for anisometric geometries that are otherwise poorly sampled using conventional diffusion-based models. Such precise control over global morphology could, in principle, enable the rapid construction of complex, higher-order protein architectures from modular protein building blocks, analogous to the programmable assembly achieved with DNA origami (*36*) - a possibility also explored through a concurrently developed CAD blueprint–guided diffusion framework (Y. Qi and T. Kortemme, personal communication).

Moreover, this framework extends to the design of functional binders within complex molecular environments. In the MD2 case, volumetric conditioning redirected sampling toward a spatially restricted, pocket-proximal region that was inefficiently sampled by hotspot-guided diffusion alone. In the case of TLR3, it enabled the precise extension of a pre-existing binder scaffold toward an adjacent ligand-binding surface while preserving the parental binding mode, thereby converting structural intuition regarding the mechanism of antagonism into designable scaffolds with validated target engagement. Together, these examples illustrate that ProteinSketch is not limited to arbitrary geometric sculpting but can serve as a practical framework for encoding biologically relevant spatial hypotheses into protein design, with particular promise for targets in which functionality depends on precise occupancy, exclusion, and interface positioning.

Several limitations of the ProteinSketch framework also point to directions vital for further development. First, ProteinSketch currently relies on RFdiffusion as a modular generative engine to translate user-specified spatial intent into protein backbones. Our volumetric benchmarks revealed a clear trade-off between geometric fidelity and designability, likely reflecting tension between the externally imposed guidance potential and RFdiffusion’s learned structural priors during denoising. Although the current framework outperformed the alternative generative approaches evaluated here, not every user-specified topology or envelope can yet be converted into a readily foldable structure. Adaptive conditioning strategies that dynamically modulate the strength of spatial guidance according to model-derived estimates of local structural plausibility may improve the balance between constraint satisfaction and biophysical feasibility. More generally, generative frameworks that incorporate topology and volumetric constraints as native conditioning inputs, rather than relying solely on externally imposed guiding potentials, may further enhance this balance. Second, from a human–computer interaction perspective, ProteinSketch requires users to construct new backbones by manipulating SSEs individually. Future interfaces could support the coordinated editing of multiple elements under explicit geometric relations, such as symmetry, alignment, and spacing, enabling a more efficient creation of larger and more complex architectures. Finally, integration with databases comprising natural protein motifs and domains could allow ProteinSketch to retrieve and present, within the VR environment, structures related to a user’s sketch, enabling designers to adapt, recombine, and graft compatible structural motifs rather than constructing every backbone *de novo*. Collectively, these advances could further extend ProteinSketch into a more capable interactive platform for human-guided, mechanism-aware design.

## Acknowledgments

This work was supported by the Korea Bio Data Station through the provision of computing resources (KBDSC-2025-KBDS-0045), and the National Supercomputing Center through the provision of supercomputing resources, including technical support (KSC-2025-CRE-0040).

## Author contributions

Conceptualization: H.M.K., S.-H.B., H.P., D.M., J.L., H.L.

Methodology: H.M.K., S.-H.B., H.P., D.M., J.L., H.L., S.-H.L., Y.G.O., T.J., S.S., S.-J.L., H.P., S.J.

Software: S.-H.B., H.P., H.M.K., D.M., H.L., S.-H.L., T.J., S.S., S.-J.L., H.P., S.J., J.L. Validation: D.M., J.L., H.L., Y.G.O., Y.-E.K.

Formal analysis: D.M., J.L., H.L., Y.G.O., H.P., S.J.

Investigation: J.L., H.L., Y.G.O., Y.-E.K., W.J., D.S.L., J.A., R.C.

Writing - original draft: H.M.K., S.-H.B., H.P., D.M., J.L., H.L., S.-H.L., Y.G.O., Y.-E.K., S.S., S.-J.L.

Writing - review & editing: H.M.K., S.-H.B., H.P., D.M., J.L., H.L., S.-H.L., Y.G.O., Y.-E.K., S.S., S.-J.L.

Visualization: H.M.K., S.-H.B., H.P., D.M., J.L., H.L., S.-H.L., Y.G.O., Y.-E.K., S.S., S.-J.L.

Supervision: H.M.K., S.-H.B., H.P.

Project administration: H.M.K., S.-H.B., H.P.

Funding acquisition: H.M.K., S.-H.B., H.P., J.L., W.J., R.C.

## Data, code, and materials availability

All data and code needed to evaluate and reproduce the conclusions in the paper are present in the paper and/or the Supplementary Materials. The atomic coordinates and cryo-EM maps have been deposited in the Protein Data Bank (PDB) and Electron Microscopy Data Bank (EMDB), respectively. The accession numbers are as follows: PDB ID 23YJ and EMD-69363 for Sphere; PDB ID 23ZT and EMD-69405 for Rod; PDB ID 23YQ and EMD-69370 for Curved arc; and PDB ID 23YT and EMD-69373 for Cone, respectively. The ProteinSketch volume potential for RFdiffusion is available at https://github.com/HParklab/ProteinSketch/. Additional protein design datasets, including experimentally validated backbone-sketching and volume-conditioned monomer and binder designs, have been deposited in Zenodo (*37*). Any other data supporting the findings of this study are available from the corresponding author upon reasonable request.

## Supplementary Materials

Figs. S1–S34

Tables S1–S8

Materials and Methods

Movies S1–S4

